# Reliability of fMRI time series: Similarity of neural processing during movie viewing

**DOI:** 10.1101/158188

**Authors:** Ralf Schmälzle, Martin A. Imhof, Clare Grall, Tobias Flaisch, Harald T. Schupp

**Affiliations:** Department of Communication, Michigan State University, USA; Department of Psychology, University of Konstanz, Germany

## Abstract

Despite its widespread use in neuroscience, the reliability of fMRI remains insufficiently understood. One powerful way to tap into aspects of fMRI reliability is via the inter-subject correlation (ISC) approach, which exposes different viewers to the same time-locked naturalistic stimulus and assesses the similarity of neural time series. Here we examined the correlations of fMRI time series from 24 participants who watched the same movie clips across three repetitions. This enabled us to examine inter-subject correlations, intra-subject correlations, and correlations between aggregated time series, which we link to the notions of inter-rater reliability, stability, and consistency. In primary visual cortex we found average pairwise inter-subject correlations of about *r* = 0.3, and intra-subject correlations of similar magnitude. Aggregation across subjects increased inter-subject (inter-group) correlations to *r* = 0.87, and additional intra-subject averaging before cross-subject aggregation yielded correlations of *r* = 0.93. Computing the same analyses for parietal (visuospatial network) and cingulate cortices (saliency network) revealed a gradient of decreasing ISC from primary visual to higher visual to post-perceptual regions. These latter regions also benefitted most from the increased reliability due to aggregation. We discuss theoretical and practical implications of this link between neural process similarity and psychometric conceptions of inter-rater reliability, stability, and internal consistency.

## Introduction

There has been an intense discussion regarding the reproducibility of research findings in functional neuroimaging (Poldrack et al., 2017) as well as science more broadly (Ioannidis, 2005; Loken and Gelman, 2017). While the debate has focused on statistical power and research practices, an important underlying topic is the reliability of the blood oxygen level dependent (BOLD) signal. Reliability concerns the precision of brain activity measures and a lack of reliability limits the validity and trustworthiness of results. Previous work on BOLD-fMRI reliability has primarily examined the temporal stability of task-related activations (Caceres, Hall, Zelaya, Williams, and Mehta, 2009; Chen and Small, 2007; Plichta et al., 2012; Specht, Willmes, Shah, and Jäncke, 2003) (Aron, Gluck, and Poldrack, 2006; Bennett and Miller, 2010; Brandt et al., 2013; Stark et al., 2004). A complementary way to tap into this issue is via the inter-subject correlation (ISC) approach. ISC analysis assesses the similarity of fMRI time series across individuals who are exposed to the same stimulus and reveals where and to what extent the continuous brain responses concur across receivers. For example, a landmark study by Hasson and colleagues showed that individuals exposed to a 30 minute excerpt from a Western movie exhibited robustly correlated time series in brain regions devoted to visual and auditory processing as well as frontal and limbic cortex (Hasson, Nir, Levy, Fuhrmann, and Malach, 2004). Subsequent research has confirmed inter-subject correlations in large scale brain networks across a broad range of dynamic, natu stimuli (Abrams et al., 2013; Chen et al., 2016; Hasson, Malach, and Heeger, 2010; Hasson et al., 2004; Imhof, Schmälzle, Renner, and Schupp, 2017; Kauppi, Jääskeläinen, Sams, and Tohka, 2010; Schmälzle, Häcker, Renner, Honey, and Schupp, 2013; Schmälzle, Häcker, Honey and Hasson, 2015).

Central to ISC analysis is the notion of similarity between functional brain responses, which relates to conceptions of reliability in measurement theory and psychology (Hasson et al., 2010). In brief, reliability refers to the quality and dependability of a measure, which encompasses different aspects of consistency and reproducibility (Shrout and Lane, 2012). One aspect of reliability is inter-rater reliability, i.e. the consensus or agreement among measures obtained from different raters. ISC analysis relates to this as it reflects the degree to which regional brain time courses from two persons who are both exposed to the same stimulus give similar ‘answers’. Another dimension of 32 reliability is temporal stability, typically assessed via test-retest measures. Although ISC analysis has been introduced to study inter-subject similarity, it can be easily adapted to look at intra-subject similarity as well: if a person views the same movie repeatedly, one can compare the similarity of the neural time courses across runs. Lastly, the ISC framework can be linked to the internal consistency aspect of reliability: Consistency refers to the extent to which different items proposed to ‘measure the same thing’ will produce similar scores. This is commonly assessed by computing the inter-item covariance and related metrics, such as Cronbach’s alpha (Davidshofer and Murphy, 2005). Importantly, the individual units that aim to measure the same construct do not have to be items, but can also refer to raters, in which case consistency blends with inter-rater reliability. This suggests that we may treat the measures from each individual as items that tap the common function of a region and aggregate across individuals to form more robust measures (e.g. the population-averaged response to a movie in limbic brain regions). Overall, there are obvious parallels between conceptions of reliability and the ISC approach, but except for one review article that touched on these issues (Hasson et al., 2010), empirical work remains limited.

This study examines fMRI responses during naturalistic viewing and assesses aspects of fMRI reliability. We do so by computing the inter-and intra-subject similarity of regional neural time-courses, and by demonstrating how aggregation of individuals into groups, comparable to aggregating items into scales, leads to very reliable measures of the aggregate regional response. Building upon previous research, the present study used movie clips, which are known to collectively engage widespread regions of posterior cortex. Each movie was repeated three times to enable test-retest analyses. The main analysis focuses on the primary visual cortex, the dorsal attention or visuospatial network anchored in the parietal cortex, and the anterior cingulate cortex, which is a central hub in the saliency network (Shirer, Ryali, Rykhlevskaia, Menon, and Greicius, 2012). In addition to these primary regions of interest, we report results for all additional regions that were expected to be engaged by the movie and are known as nodes of major brain networks related to vision, dorsal attention, saliency, executive control, and default mode processing (Shirer et al., 2012). Based on theoretical frameworks suggesting an information flow from primary to higher-order visual association areas, to multimodal paralimbic regions (Fuster, 2003; Mesulam, 1998; Pandya and Yeterian, 2003), we expected to see a gradient of high inter-subject correlations in primary visual regions, to medium strength in parietal regions, to lower, but still positively correlated responses in anterior cingulate cortex.

## Methods

### Participants

Twenty-four healthy adults (*mean age* = **23.2** years, **SD** = 2.88; range = 20 to 36 years, 13 females) with normal or corrected-to normal vision and no history of neurological of psychiatric disease participated in this study. One additional participant was immediately replaced due to scanner artifacts. All participants provided written consent to the study protocol, which was approved by the local ethics committee. Participants received either course credit or monetary compensation.

### Stimulus and Procedure

The movie clips used in this study were extracted from 75 Hollywood feature films and TV broadcasts. They consisted of positive, erotic materials depicting partly or completely naked heterosexual couples engaging in sexual activity, but they were not explicitly pornographic. The length of the erotic movie stimulus was 4 minutes and 30 seconds, comprising six concatenated short clips. The movies were shown on a MR-compatible visor system (VisualSystem, NordicNeuroLab, Inc.) with a screen resolution of 800 x 600 pixels, and timing was time-locked to the scanner triggers using Presentation software (Neurobehavioral Systems, Inc.). Participants were instructed to freely watch the clips while holding their heads still. Participants also viewed an assortment of neutral movie clips, but comparison of responses between the erotic and neutral movies is beyond the scope of this study and will be reported elsewhere. However, analysis of the neutral clips shows similar findings across levels of aggregation as reported here. Participants saw each movie three times in alternating order, randomly starting with either the positive or the neutral video assortment. After the viewing task we acquired a high-resolution structural scan from each participant.

### MRI acquisition and analysis

MRI data were acquired using a Philips Intera 1.5 90 Tesla scanner equipped with Power Gradients. BOLD data was measured using a Fast Echo Planar Imaging sequence (FFE-EPI, T2*-weighted, 90 degree flip angle, TR = 92 2500 ms, TE = 40 ms, ascending-interleaved slice order, in plane resolution of 3 x 3 mm, slice thickness of 3.5 mm, 32 slices, no gap, FOV = 240 x 240 mm). We obtained 110 functional volumes during each presentation. Structural images were obtained at the end of the experiment using a standard T1-weighted high resolution scan with a voxel size of 1 x 1 x 1 mm (T1TFE, FOV = 256 x 256 mm, 200 sagittal slices). Field of View was adjusted in line with the AC-PC plane.

### FMRI Preprocessing and Data Extraction

Data was preprocessed using SPM12 for realignment, slice time correction, and DARTEL normalization to the IXI500 template (http://brain-development.org/ixi-dataset/). Further processing was carried out in Python 2.7 using the nilearn package (Abraham et al., 2014) and custom code. Specifically, functional time courses we e extracted from regions provided by the functional atlas of Shirer and colleagues (Shirer et al., 2012) (see Supplementary Figure s1 for an overview). We chose to focus on regions rather than individual voxels because regions are a common, established measurement unit in neuroimaging, and because region-level as opposed to voxel-level responses should be less sensitive to anatomical differences and allow for easier communication of results at the meso-scale (i.e. less than 100 regions vs. more than 40.000 voxels). However, the same ideas can be applied to voxel-level-analysis. In addition to the three primary regions of interest, i.e. primary visual, parietal, and anterior cingulate cortex, we also examine the results for additional regions that might be expected to be engaged by the visual stimulus (e.g. higher visual system, executive control network, default mode, see Supplementary Materials) as well as the left and right auditory cortex as control regions. Furthermore, we added subcortical regions for exploratory purposes: the bilateral thalamus, the amygdala (Tzourio-Mazoyer et al., 2002), and the ventral striatum (Tziortzi et al., 2014). FMRI time courses were extracted from these regions and data were filtered (0.01 - 0.12 Hz), detrended, and motion parameters were added as confounds. The first 10 volumes that might be affected by signal transients were removed, and the time courses were z-scored. Thus, we obtained a neural time course consisting of 100 samples for each of the 38 regions, from 24 participants and 3 repetitions, yielding a 100 x 38 x 3 x 24 data matrix that served as the basis for further analysis.

### Data analysis

For each region, we computed Pearson correlations between the fMRI time series to capture i) the similarity of individual regional time courses between participants (inter-subject correlation), ii) the similarity of time courses from the same participant over repeated viewings (intra-subject correlation), and iii) the similarity of time courses after initial averaging over multiple participants (inter-subject correlations of aggregate time-courses) (Hasson et al., 2004). For the inter-subject correlation analysis (i) we begin with the fMRI time series recorded from 24 viewers during the first viewing of the movie clips. To measure the similarity of neural responses in each region, we compute the correlation between time-series from a given region (e.g., *r_S1V1(subject1-1st viewing)-vs.-S2V1(subject2-1st viewing)_*. This analysis is then computed for all pairs of participants (i.e. ((24 *24) -24)/2 = 276 values), and the resulting values are Fisher-*z*-transformed, averaged, and finally re-transformed to an *r*-value that represents the average similarity of time series across viewers in a given region. For the intra-subject analysis (ii), we proceed in an analogous manner, but compute correlations between time courses from the same individual across the three viewings of the same movie (i.e., *r_S1V1-vs.S1V2_, r_S1V1-vs.-S1V3_, r_S1V2-vs.S1V3_*, etc.). To compare intra-subject correlations with inter-subject correlations, we also computed inter-subject correlations also across different runs (e.g., *r_S1V1-vs.-S2V2_*) to account for potential order effects. Finally, for the aggregated inter-subject, or inter-group analyses (iii), we average the time series from multiple viewers (e.g. from pairs, triplets, etc.) to create more reliable signals, and then assess the correlation of aggregated signals at progressive levels of aggregation. For example, we begin by taking the average time series from visual cortex for viewers A and B, and correlate this average time series against the average of neural time series from viewers C and D. Because there are multiple ways to split up the viewers into groups of two, we compute these inter-group correlations for 1000 permutations of the grouping process. This scheme is applied from average pairwise, or dyadic ISC up to the *split-half* ISC, which represents the correlation between the average time series from one half of the group against the average of the remaining half.

## Results

### Inter-subject correlations

We begin by examining the inter-subject correlations of neural time-series between individual viewers in the visual cortex (Figure 1A - left panel). Figure 1D shows the neural time courses from two random participants' visual cortex during the first viewing. As can be seen, viewing the same time-locked movie sequences prompts similar time courses and the inter-subject correlation for this random pair of subjects amounts to *r_S1V1vsS2V1_* = 0.22. Across all pairwise inter-subject correlations, we find an average *r_interSC: pairwise_* = 0.35. Next, we start aggregating time courses of individual viewers. As a first aggregation step, we sample two random viewers, average their visual cortex time course, and compare this 2-viewer-average time course against another 2-viewer average time course from another random pair (*e.g. r_(avg(S1V1, S2V1)-vs.-avg(S3V1,S4V1))_*).Figure 1D shows the change in correlations for example time courses averaged over two participants. Figure 1D (bottom left) expands this procedure to averages of 6 persons each (*e.g. r_avg(S1,S2,S3,S4,S5,S6)-vs.-avg(S7,S8,S9,S10,S11,S12))_*). As expected, averaged time courses are more similar than the individual time series and the corresponding correlations increase to an average *r_interSC: 2-person averages_* = 0.52 and *r_interSC: 6-person averages_* = 0.76 across permutations. The highest aggregation step is the one that splits the dataset into two halves and measures the correlation between time course averages from two groups of twelve viewers each. As shown in Figure 1D (bottom right), these split-half time courses are highly similar with an average correlation of *r_interSC: split-half_* = 0.87 across permutations. Figure 1G generalizes this procedure to all aggregation levels and shows the evolution of ISC in primary visual cortex as we aggregate from pairwise analyses, to duplets, triplets, and all the way up to split-half correlations.

**Figure 1.**
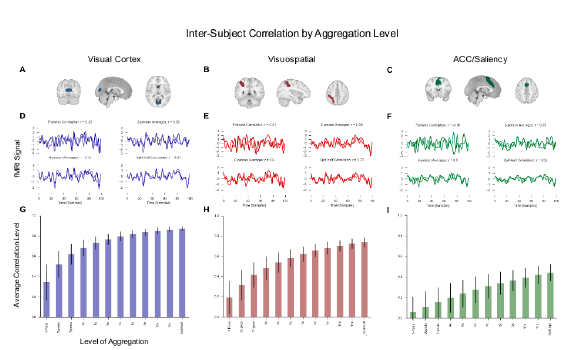
Inter-subject correlation by aggregation level for regions involved in visual, visuospatial, and saliency processing. A-C: The top panels of each subfigure show the anatomical location of the nodes (Shirer et al., 2012). D-F: The time series plot in the middle panel shows example time courses from two random viewers (top left), the averaged time courses from two viewers each (top right - first step of aggregation), the averaged time course across six viewers each (bottom left), and finally the averaged time courses of twelve vs. twelve viewers (bottom right: split-half correlation). G-I: The bar plots in the lowest panel illustrate the evolution of regional ISC across aggregation steps.

The same analytical steps as described for the visual cortex were also carried out for the signals measured in the parietal and cingulate cortex (visuospatial and saliency network). As can be seen from the corresponding plots in Figure 1E and 1H, ISC in these regions is lower compared to the visual cortex, but still detectable to the human eye. For example, the average correlation of individual time series from the visuo-spatial network is about *r_interSC: pairwise_* = 0.2, and these correlations increase up to a level of *r_interSC: split-half_* = 0.74 through averaging.

Finally, for the saliency network, individual time series exhibit relatively weak correlations around *r_interSC: pairwise_* 0.1 (Figure 1I, first bar). However, as can be seen in Figure 1F for the example time courses at the dyadic level, we sometimes find even negative values, suggesting relatively high noise. However, aggregation across individual viewers leads to progressively higher values up to a level of rinterSC: split-half = 0.44 for the correlation of two 12-person-averaged time series.

**Figure 2.**
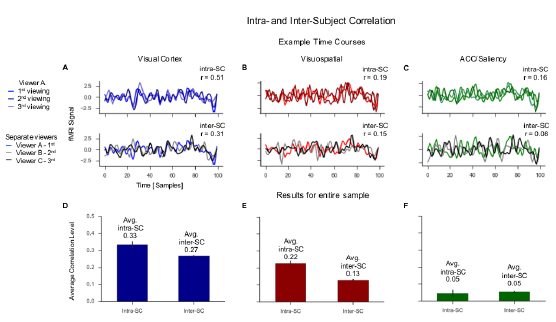
Intra-and inter-subject correlations. A-C. The top panels of the time series plots show data from one viewer across the three viewings, the lower panels show data from a different viewer for each viewing. D-F. The bar plots show results across all viewers. Intra-subject correlations are nominally higher for the visual and visuospatial network, but of similar magnitude for the saliency network.

### Intra-subject correlations

Participants in this study viewed three repetitions of the same movie clips. This allowed us to assess the similarity of regional fMRI time courses from a given viewer over repeated viewings. These intra-subject correlations can be linked to test-retest reliability, or measurement stability analysis. As can be seen in Figure 2 for one viewer, the visual-cortex time-series from different repetitions resembled each other and exhibited an intra-subject correlation of *r_intraSC_* = 0.51. Moving beyond the selected exemplar viewer, we carried out this intra-subject correlation analysis for all viewers and averaged the results: For the visual cortex we find average intra-subject correlations of *r_intraSC_* = 0.33 across viewers. This value is nominally higher than the corresponding inter-subject correlations, which amount to about rinterSC: pairwise = 0.27. Of note, because intra-subject correlations necessarily compare data from different viewings, we also computed inter-subject correlations across viewings (e.g. viewer A, 1st viewing vs. viewer B, 2nd viewing) to make the measures comparable. Therefore, inter-SC values are slightly different from the results in Figure 1, which reflected the inter-SC during the first viewing only.

A similar pattern emerges for the visuospatial network, which again yields somewhat lower similarities compared to the visual cortex. As shown in Figure 2B and E, intra-SC are somewhat higher than inter-SC. Finally, for the ACC node of the saliency network (Figure 2 C,F) we find small intra-and inter-subject correlations. Supplementary table S1 presents results for all other regions.

**Figure 3.**
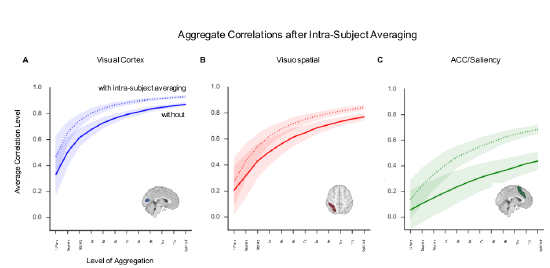
Inter-subject correlation by aggregation level with and without previous intra-subject averaging. The x-axis represents the level of data aggregation, ranging from pairwise time course correlations to split-half correlations. To account for randomness of the group-averaging process, all analyses are fully permuted and the results are averaged to produce point estimates for each step. Shaded areas indicate +/- 1 SD of the permuted ISC results

### Increasing inter-subject similarities through initial 210 within-person averaging

The presence of substantial intra-subject correlations suggests that intra-subject averaging can increase the signal to noise ratio of the time series and thus provide a more robust basis for subsequent inter-subject or inter-group correlation computations. The enhancement of ISC due to this initial within-subject averaging is illustrated in Figure 3. For comparison, this figure also includes the corresponding inter-SC values without previous intra-subject averaging (cf. Figure 1C, F, I). For the visual cortex, the pairwise inter-subject correlation increases from *r_intraSC: pairwise_* = 0.35 for single-viewing to *r* = 0.48 across the intra-subject averages from three viewings. At the level of split-half aggregation, which exhibited a split-half correlation during first viewing of *r_interSC: split half_* = 0.87, this intra-subject averaging increases the split-half reliability to *r* = 0.93. Similarly, correlations in the visuospatial network rise from *r_isc: split half_* = 0.74 without intra-subject averaging to *r* = 0.82, and for the dACC rise from *r_isc: split half_* = 0.51 without intra-subject averaging to r = 0.69.

The previous analyses were carried out on three regions selected based on their location in the processing stream (Mesulam, 1998). To provide a broader picture, we carried out the same analyses for all other 35 nodes. The results, shown in Figure 4, parallel and expand the findings for the primary visual, visuospatial, and ACC regions. In particular, we observed a decreasing gradient of ISC as we move away from visual sensory regions, and there are strong effects of aggregation. As expected for the silent-movie stimulus, the auditory cortex showed only weakly correlated responses across participants. Also, amygdala, the striatum, and some regions of the default and executive control network exhibit low correlations, suggesting that the selected experimental movie did not engage activity in these regions very consistently across viewers.

**Figure 4.**
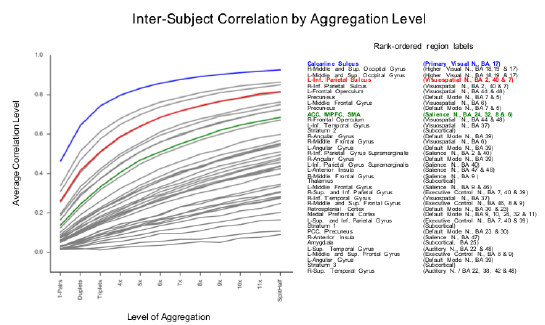
Inter-subject correlation by aggregation level (after previous intra-subject averaging). Text labels for the regions are ranked based on the level of ISC, ranging from highest in primary visual cortex to lowest in auditory cortex (See Supplementary Materials for full list and data). R = Right, L = Left, N = Network, Inf.= Inferior, Sup. = Superior.

## Discussion

The present study examined how similarly the brains of 24 participants respond during a natural movie stimulus. Similarity of neural processing was quantified via inter-and intra-subject correlation analysis. As expected, we found inter-SC to be maximal in visual cortex and lower, but still reliable across widespread brain regions implicated in higher order processing. Intra-subject correlations were generally higher than inter-subject correlations, and inter-subject averaging as well as intra-subject averaging over repeated exposures markedly increases this similarity, particularly for regions in which similarity was not evident in pairwise correlations. These inter-and intra-subjective process similarities can be linked to inter-rater, consistency, and stability aspects of reliability, and they provide important information about the reliability of functional neuroimaging

### Inter-subject correlations

The standard, pairwise inter-subject correlation analysis confirms correlated responses in visual sensory, perceptual, and higher order regions during movie viewing. For the visual cortex, we find a robust inter-subject correlation of *r_intraSC: pairwise_* = 0.35 on average between the time courses from different viewers. Many other regions beyond primary visual cortex also exhibit similar processing across viewers, such as the highlighted visuospatial region, *r_intraSC: pairwise_* = 0.2, along with other visual associative regions (see Table s1). The ACC, however, exhibits less strong correlations - ranging only around 0.1 at the pairwise level. This pattern of results aligns with previous ISC findings (Hasson et al., 2004; Jääskeläinen et al., 2016) and is compatible with models of neural information processing that propose a gradient of conservation from primary to associative memory structures (Fuster, 2003; Mesulam, 1998; Pandya and Yeterian, 2003). These models, suggest the ACC as a region that integrates incoming external information with internal demands pertaining to motivation, emotion, and homeostasis, which could explain why it exhibits lower correlations across and within viewers. The varying strength of ISC in sensory, perceptual, and post-perceptual regions underscores that post-sensory regions do not just respond in a reflexive, stimulus driven manner, but rather engage with the incoming stimulus based on the match between the stimulus and regional response sensitivities. In this sense, a movie provides each brain region with a time-varying landscape of action-opportunities (Gibson, 1977), and the resulting correlations of fMRI responses point to commonalities in how perceptual and cognitive-emotional systems navigate this landscape.

Based on this reasoning, we suggest that inter-SC can be understood as a neural counterpart of inter-rater agreement (Krippendorff, 2004; Riff, Lacy, and Fico, 2014) at the level of individual brain regions, and that inter-SC of neural processes can quantify the extent to which a movie, or any other communicative signal, is successfully transmitted into a recipient's brain (Hasson, Ghazanfar, Galantucci, Garrod, and Keysers, 2012). This naturally leads to questions about the content or nature of these reliably shared responses. While acknowledging the limits of reverse inference (Poldrack, 2006), the question seems at least partially addressable for sensory and perceptual regions: By parametrizing the movie into constituent features, one can identify how these are tracked by regional responses or can reverse-correlate from neural responses back to movie features (Naselaris, Kay, Nishimoto, and Gallant, 2011; Ringach and Shapley, 2004). For higher-order regions, however, which are less tightly coupled to immediate stimulus properties and less reliably correlated across receivers, our knowledge about the psychological interpretation of activity remains limited (e.g. dACC or default mode regions). However, inter-subject correlation methods provide unique opportunities to examine these higher-order integrative processes, for instance, by systematically varying the psychological state across or within individuals.

### Intra-subject correlations

Intra-subject correlations assess the stability and individual variability of movie-evoked responses. Overall, we find intra-subject correlations to be slightly higher than inter-subject correlations whenever there is reliable inter-or intra-SC in the first place (e.g. for 15 out of 15 regions with pairwise values above 0.1). This is noteworthy for two reasons: First, we analyzed signals from relatively large regions, which should have reduced the impact of anatomical differences that might favor higher intra-subject correlations, but still found higher intra-than inter-subject responses. This points to individual differences in functional brain responses, which are a topic of ongoing research (Campbell et al., 2015; Finn et al., 2017). Second, although slightly higher, the intra-subject correlations were generally of a comparable magnitude to inter-SC (i.e. both around *r* = 0.3 in visual cortex). This value range may at first seem low compared to what is considered adequate in psychometrics, but we note that the underlying data are based on one single viewing. As such, the intra-subject correlations could be substantially increased by averaging across multiple trials - just as for inter-subject correlations, or classical task-based studies that report higher stability (Plichta et al., 2012;). A further noteworthy finding from the intra-subject analysis concerns the dACC, where intra-SCs were not even nominally higher than inter-SC (see Figure 2 right panel). This suggests pronounced intra-individual variability (e.g. motivation, attention, habituation) that is apparently comparable in size to inter-individual differences, and poses a challenge for reliable measurement (Nord, Gray, Charpentier, Robinson, and Roiser, 2017).

One key issue for intra-subject analyses is that the underlying phenomenon might vary over time. Indeed, viewers who watched the erotic clips the second time might have started to either habituate or sensitize, or form predictions. This would likely change motivational salience and cognitive control processing and thus affect intra-subject correlations, which certainly cannot result from anatomical differences. Overall, more work is needed to examine intra-subject correlations, and only few studies have begun to look more deeply into these issues, which can complement large population studies (Savoy, 2006).

### Averaging across viewers to create more reliable aggregate measures

Finally, we examined the effects of aggregation on fMRI time series correlations and showed that this yields highly reliable aggregate measures. In psychometrics, it is commonplace to combine individual items that tap into a common construct into a scale. Importantly, the idea of aggregating over multiple items can be applied when raters are treated as items, which effectively blends the notion of internal consistency and inter-rater reliability. As shown in Figures 1 and 3, the similarity between averaged time courses increases markedly as we ‘average in’ data from additional viewers. Thus, high inter-group correlations emerge for many regions for which inter-subject correlations were barely detectable. For example, the high correlation around r = 0.9 for the visual cortex shows that this procedure drastically reduces noise due to technical factors like scanner noise and non-shared signals such as individual differences.

One important benefit of this boost in reliability is related to the fact that ‘reliability limits validity’, i.e. that the correlation between two measures will, on average, not exceed the product of their individual reliabilities. Consider, for example, a scenario in which the goal is to examine fMRI-and fNIRS responses during movie processing (Hasson et al., 2008). In such cases, group-averaging can markedly improve the accuracy at which similarities between the measures can be assessed and thereby aid methods comparison. Perhaps more importantly, higher reliability will also improve correlations between fMRI measures and psychological variables, such as continuous attention (Nummenmaa et al., 2014), or psychological traits and other external variables (Berkman and Falk, 2013; Cohen, 1992; Poldrack et al., 2017).

Beyond better measurement, however, the issue of aggregation across individuals also raises interesting theoretical questions in their own right. If a region has no common signal to aggregate, this suggests that it is either not engaged by the stimulus in the first place (e.g., olfactory cortex during visual stimulation), that the region was not scanned reliably, or that functional anatomical or psychological factors produce variability between viewers. This latter point, together with the above discussion of intra-subject analysis, speaks to long-standing distinctions between commonality and individual differences in psychology (Lamiell, 2003). In this sense, the benefit of aggregation is that we ‘average in’ common signal and obtain highly reliable measures, but at the same time we ‘average out’ idiosyncratic information about anatomical or psychological differences; whether inter-or intra-SC analyses are to be chosen thus depends on the specific research question. Of note, this is not only an issue for aggregated ISC analyses, but is actually implicit in second level tests in the widespread statistical parametric mapping approach (Friston et al., 1994).

### Implications for reliability of fMRI, limitations, and questions for future research

Studying how continuous and dynamic stimuli engage the brain is a relatively young field and thus a number of open questions remain. Obviously, the current study is limited to visual processing and to the specifics of the movie clips at hand. However, the observed findings regarding the pattern of reliability replicates when we perform the same analyses for a different visual movie (neutral clips) and there is first evidence that this also applies to other modalities (e.g. (Lerner, Honey, Silbert, and Hasson, 2011). Second, we decided to study correlations between regional responses rather than individual voxels. The reason for this is that we are primarily interested in the level of meso-scale brain systems linked to affect and attention, which are plausible targets for media effects. Additionally, examining regional responses helps to overcome anatomical differences at smaller scales and provides a straightforward, scalable strategy. Of course, much can be learned from studying voxel level similarities or using these for functional normalization as in Hyperalignment (Haxby et al., 2011) or shared response models (Chen et al., 2016). Third, we communicate relevant information about variability via the error bars in Figures 1–figures 3, but otherwise avoid sole reliance on null-hypothesis tests. Conventional significance tests are not immediately relevant to our main points regarding inter-SC, intra-SC, their spatial distribution, and the effects of aggregation (Gigerenzer, 2004), and different types of tests with different sample sizes would be adequate for inter-SC, intra-SC, and inter-group similarity. For example, if the goal is to test whether a distribution of pairwise ISCs is significantly different from zero, then recent work by Chen and colleagues (2016) provides a discussion of the statistical dependency among pairwise correlations. If one wants to test whether two individual time series - either from two viewers, viewings, or two aggregate time-series - are significantly related, then classical time-series methods are appropriate (Cochrane and Orcutt, 1949; Hamilton, 1994). Fourth, we present findings from the perspective of classical test theory (CTT) because it is intuitive and most researchers in neuroscience and psychology are familiar with it. However, generalizability theory (Cronbach, 1972; Gao and Harris, 2012) offers an advanced framework to decompose different facets of variation (e.g. persons, items, raters, time, or setting) and assess their relative contribution. Future multi-site, multi-stimulus, multi-method, and population-level initiatives may perform G analyses to produce comprehensive neural reliability maps for different facets (Dubois and Adolphs, 2016; O‘Connor et al., 2016). Related to this, the time-series correlations we report are linked and in some cases mathematically equivalent to intra-class correlations (ICC) (Shrout and Lane, 2012). Finally, it will be interesting to expand the notion of process similarity beyond relationships among corresponding single brain regions: well-controlled movies provide an ideal tool to study similarities of dynamic functional connectivity (Bassett et al., 2011; Simony et al., 2016), or similarities of more complex network measures (Andric, Goldin-Meadow, Small, and Hasson, 2016; Pannunzi et al., 2016; Sizemore, Giusti, Betzel, and Bassett, 2016; Wang et al., 2017). Critically, these findings should not be taken to evaluate the reliability of fMRI as a whole. There is no such thing as the *reliability* of a measure and a multitude of factors will affect the magnitude of correlations. For example, the relatively modest ISC in ACC should not be taken as universal, as different stimuli (e.g a movie with shorter duration or strong negative contents), different participants (e.g. patients), or different scanning parameters could change the magnitude of the correlations. However, the outlined conceptual framework is robust to such specifics and disentangling different facets of within and between person variation can contribute to the development of fMRI in the next decades (Dubois and Adolphs, 2016).

## Summary and conclusion

This study examined inter-and intra-subject correlations during movie viewing and linked the underlying notion of neural process similarity to aspects of reliability. Especially in the era of multi-lab population neuroimaging initiatives, this approach holds great promise for probing a wider psychological repertoire in a highly reliable and controlled, but eminently scalable way (O’Connor et al., 2016). This may lead to neurometric databases of functional brain responses to movies or stories and yield important information about the distribution of normal brain function and for the diagnosis of disorders (Dubois and Adolphs, 2016). Importantly, while this paper focuses on methodological concepts, the measures of neural process similarity are of interest in their own right and offer new means to examine inter-and intra-individual neural differences in higher-order cognitive and motivational processes (Campbell et al., 2015; Hasson et al., 2009; Honey, Thompson, Lerner, and Hasson, 2012; Imhof et al., 2017; Naci, Cusack, Anello, and Owen, 2014; Schmälzle et al., 2013).

## Acknowledgements

We thank the developers of NiLearn, Nibabel, and Seaborn for their contributions to open source software. H.T.S. and M.A.I. were supported in part by the German Research Foundation [DFG, FOR 2374].

## Supporting Information

Supplemental material for this article is available at *https://github.com/nomcomm/NeuralProcessReliability*. This repository contains code that was used to produce the results and figures in the paper

## Supplementary Information

**Supplementary Figure 1.**
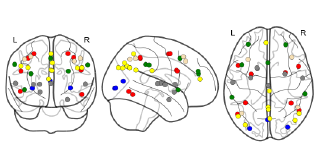
Center coordinates of regions used for extraction of neural time series. Colors indicate membership in large-scale networks, blue - Visual, red - Dorsal Attention, green - Saliency, yellow - Default Mode, wheat - Executive Control, gray nodes added for exploratory purposes.

**Supplementary Table 1.** Regional metrics for inter-subject correlation, aggregate-level inter-subject correlations, and intra-subject (test-retest) correlations. The first column displays the average correlation value across all inter-subject correlation pairs (*r_S1V1-vs.-S2V2, rS1V1-vs.-S3V1_, … r_S2V13-vs.-S24V1_*). The second column shows a variant of the first column in which pairwise correlations are computed across runs (e.g. *r_S1V1-vs.-S2V2_*). The third column shows the average intra-subject correlation between the first and second viewing. The last column shows the split-half correlation, i.e. intersubject or inter-group correlations between the averaged time course from 12 viewers vs. the averaged time course of the remaining 12 viewers (averaged across permutations of the group-forming process).

|  | *Pairwise inter-subject correlation* | *Pairwise inter-subject correlation (1st vs. 2nd viewing)* | *Intra-subject correlation (1st vs. 2nd viewing)* | *Split-half correlation* |
| --- | --- | --- | --- | --- |
| *Prim_Visual_01* | 0.35 | 0.31 | 0.41 | 0.87 |
| *High_Visual_01* | 0.19 | 0.16 | 0.26 | 0.74 |
| *High_Visual_02* | 0.24 | 0.18 | 0.26 | 0.79 |
| *Visuospatial_01* | 0.07 | 0.09 | 0.08 | 0.5 |
| *Visuospatial_05* | 0.06 | 0.05 | 0.07 | 0.42 |
| *Visuospatial_02* | 0.2 | 0.13 | 0.23 | 0.74 |
| *Visuospatial_06* | 0.18 | 0.16 | 0.25 | 0.74 |
| *Visuospatial_07* | 0.08 | 0.06 | 0.13 | 0.52 |
| *Visuospatial_03* | 0.14 | 0.11 | 0.19 | 0.66 |
| *Visuospatial_04* | 0.09 | 0.05 | 0.14 | 0.52 |
| *Visuospatial_08_8* | 0.01 | 0.02 | 0.01 | 0.17 |
| *Anterior_Salience_01* | 0.01 | 0.03 | 0.03 | 0.12 |
| *Anterior_Salience_02* | 0.01 | 0.02 | 0.02 | 0.18 |
| *Anterior_Salience_03* | 0.06 | 0.06 | 0.02 | 0.44 |
| *Anterior_Salience_04* | 0.04 | 0.02 | 0.02 | 0.34 |
| *Anterior_Salience_05* | -0.01 | 0.01 | 0.06 | -0.04 |
| *Posterior_Salience_02* | 0.06 | 0.06 | 0.17 | 0.43 |
| *Posterior_Salience_06* | 0.04 | 0.04 | 0.1 | 0.35 |
| *LECN_01* | 0 | 0.01 | 0.05 | 0.05 |
| *LECN_03* | 0.04 | 0.03 | 0.06 | 0.31 |
| *RECN_01* | 0.05 | 0.02 | 0.11 | 0.39 |
| *RECN_03* | 0.06 | 0.03 | 0.13 | 0.37 |
| *Dorsal_DMN_01* | 0.04 | 0.03 | 0 | 0.31 |
| *Ventral_DMN_04* | 0.03 | 0.02 | 0.14 | 0.23 |
| *Ventral_DMN_09* | 0.06 | 0.03 | 0.14 | 0.41 |
| *Dorsal_DMN_02* | 0.05 | 0.03 | 0.14 | 0.34 |
| *Dorsal_DMN_06* | 0.01 | 0 | 0.02 | 0.02 |
| *Dorsal_DMN_04* | 0.02 | 0.01 | 0.01 | 0.26 |
| *Precuneus_DMN_01* | 0.03 | 0.03 | 0.06 | 0.28 |
| *Precuneus_DMN_02* | 0.08 | 0.09 | 0.1 | 0.51 |
| *Ventral_DMN_06* | 0.12 | 0.07 | 0.15 | 0.63 |
| *Auditory_01* | 0.02 | 0.01 | 0.08 | 0.23 |
| *Auditory_02* | 0.02 | 0 | 0.02 | 0.15 |
| *Striatum_1* | 0.01 | 0.01 | 0.02 | 0.07 |
| *Striatum_2* | 0.03 | 0.04 | 0.01 | 0.25 |
| *Striatum_3* | 0 | 0.01 | 0.03 | -0.03 |
| *Amygdala* | 0.03 | 0.02 | 0.06 | 0.28 |
| *Thalamus* | 0.07 | 0.02 | 0.06 | 0.45 |

